# A phospholipid-dependent PDK1-AGC kinase cascade regulates pollen tube growth

**DOI:** 10.64898/2026.03.03.709238

**Authors:** Tong Zhao, Remko Offringa

## Abstract

3-PHOSPHOINOSITIDE-DEPENDENT PROTEIN KINASE1 (PDK1), a conserved master regulator of AGC kinases, is encoded by two redundant genes in *Arabidopsis thaliana, PDK1* and *PDK2. pdk1 pdk2* mutants exhibit a broad range of defects, including apolar or arrested pollen tube growth, a phenotype also observed in *agc1*.*5 agc1*.*7* mutants. Pollen-specific expression of constitutively active AGC1.5 in *pdk1 pdk2* restores polar pollen tube growth, indicating that PDK1 functions upstream of redundant AGC1.5/AGC1.7 signaling in this process. In contrast, the *PDK1* splice variant PDK1S0, lacking the phospholipid-binding PH domain, cannot restore polar pollen tube growth. Our results indicate a key role for the phospholipid PI(4,5)P_2_ in recruiting PDK1 through its PH domain to establish polar pollen tube growth, as PI(4,5)P_2_ marks the pollen germination initiation site together with PDK1, it forms a dome at the plasma-membrane of the pollen tube tip beneath which PDK1 remains largely cytosolic and exhibits reciprocal feedback regulation with the PDK1-AGC1.5/1.7 kinases. Defects in endocytosis and actin organization further support that phospholipid-dependent PDK1–AGC signaling maintains pollen tube growth polarity.

## INTRODUCTION

Pollen tube growth, a characteristic form of tip growth, is essential for the double fertilization process in flowering plants (Higashiyama and Yang, 2016). The regulation of pollen tube growth involves multiple signaling pathways and cellular processes.

Phosphatidylinositol 4,5-bisphosphate (PI(4,5)P_2_), which localizes to the apical region of the pollen tubes, plays an important role in polarized tube growth (Heilmann and Ischebeck, 2016). PI(4,5)P_2_ is synthesized by PHOSPHATIDYLINOSITOL-4-PHOSPHATE 5-KINASES (PIP5Ks), and since five of the encoding genes in *Arabidopsis thaliana* (Arabidopsis) act redundantly in pollen, *pip5k* loss-of-function mutants either show minor defects in pollen tube growth or are defective in pollen germination(Ischebeck et al., 2008; Sousa et al., 2008; Zhao et al., 2025). In contrast, *PIP5K* overexpression results in branched or stunted pollen tube growth (Ischebeck *et al*., 2008; Sousa *et al*., 2008; Zhao et al., 2010). PI(4,5)P_2_ is crucial for endocytosis and actin dynamics. Downregulation of *PIP5K6* expression reduces endocytosis (Zhao *et al*., 2010); whereas overexpression of *PIP5K10* or *PIP5K11* disrupts actin organization in pollen tubes(Ischebeck et al., 2011).

Disrupting fine actin filaments or mutating various *ACTIN BINDIG PROTEIN* (*ABP*) genes results in curly, swollen, or stunted pollen tubes (Cheung et al., 2010; Qu et al., 2013; Ye et al., 2009; Zheng et al., 2013). Actin dynamics and organization, in turn, influence vesicle trafficking and endo/exocytosis (Feiguelman et al., 2018; Scheible and McCubbin, 2019).

AGC kinases are a group of conserved protein serine/threonine kinases in eukaryotes that participate in a broad spectrum of biological processes, including regulating pollen tube growth in plants (Rademacher and Offringa, 2012; Zhang and McCormick, 2009). The 3-PHOSPHOINOSITIDE-DEPENDENT PROTEIN KINASE1 (PDK1) functions as a master regulator of other AGC kinases in eukaryotic cells (Mora et al., 2004; Tan et al., 2020; Voordeckers et al., 2011; Xiao and Offringa, 2020). Besides its kinase domain, PDK1 possesses a PH domain that, upon binding to phospholipids, alleviates its autoinhibitory effect on the kinase domain while simultaneously destabilizing the protein as a feedback control mechanism (Bayascas et al., 2008; Levina et al., 2022). Arabidopsis possesses two PDK1-encoding genes, *PDK1* and *PDK2*, whose double loss-of-function mutants display non-polar or arrested pollen tube growth (Xiao and Offringa, 2020). *agc1*.*5 agc1*.*7* double mutants exhibit similar pollen tube growth defects, and AGC1.5/1.7 are expressed in pollen (Zhang et al., 2009).and are phosphorylated by PDK1 *in vitro* (Zegzouti et al., 2006). Moreover, RopGEFs are phosphorylated by AGC1.5, controlling pollen tube polarity (Li et al., 2018).

Here we studied the role of PDK1 in regulating polarized pollen tube growth in more detail. We show that the non-polar or arrested pollen tube growth of *pdk1 pdk2* is rescued by pollen-specific expression of a constitutively active version of AGC1.5 (AGC1.5SD), indicating that the redundant AGC1.5 /1.7 kinases act downstream of PDK1. In contrast, the twisted pollen tube phenotype is not rescued by the *PDK1* splice variant PDK1S0, indicating that the lacking phospholipid-binding PH domain is essential for growth polarity. Our results indicate a key role for the phospholipid PI(4,5)P_2_ in recruiting PDK1 through its PH domain to establish polar pollen tube growth: PI(4,5)P_2_ marks the pollen germination initiation site together with PDK1, it forms a dome at the plasma-membrane of the pollen tube tip beneath which PDK1 remains largely cytosolic and exhibits reciprocal feedback regulation with the PDK1-AGC1.5/1.7 kinases. Finally, our results show that the phospholipid-dependent PDK1-AGC1.5/1.7 cascade guides polar pollen tube growth by regulating endocytosis and actin organization.

## RESULTS

### Polar pollen tube growth requires PH domain-dependent PDK1 activity

Previously, we reported that *pdk1-13 pdk2-4* and *pdk1-14 pdk2-4* double loss-of-function mutants (hereafter referred to as *pdk1 pdk2*) show a plethora of developmental defects (Xiao and Offringa, 2020). Most defects except for the shorter siliques were rescued by expressing splice variant *PDK1S0*, which encodes a PDK1 version lacking the PH domain (Fig. 1A). The shorter silique phenotype was also observed for the *pdk1-b pdk2-1* double mutant (Camehl et al., 2011), where *pdk2-1* is a loss-of-function allele but *pdk1-b* encodes a PDK1 version lacking the PH domain (Fig. 1A) (Xiao and Offringa, 2020). Genetic analysis indicated that the shorter siliques are caused by defects in the male gametophyte, among which defective pollen tube growth (Xiao and Offringa, 2020).

**Figure 1.**
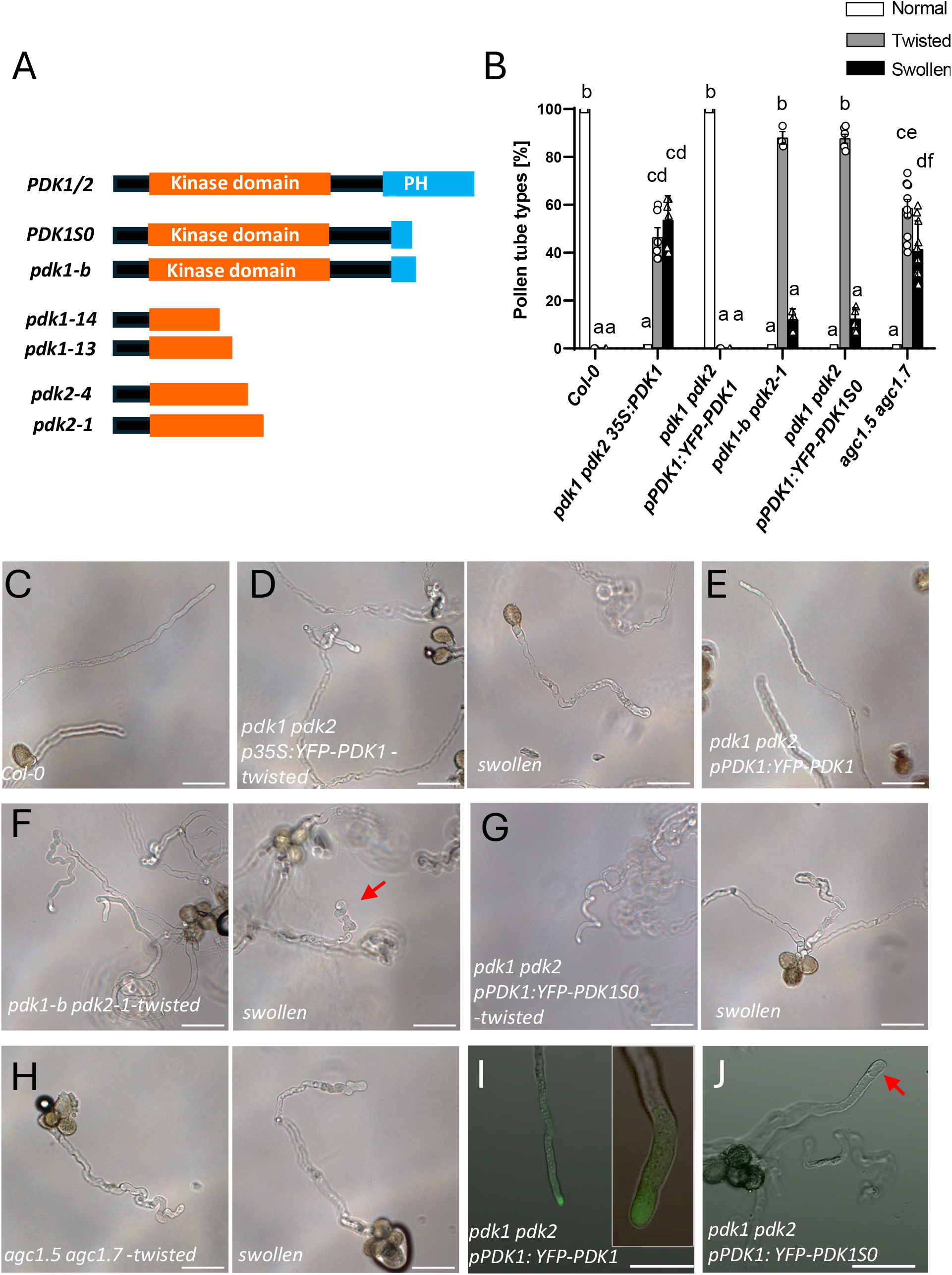
PDK1 and AGC1.5/AGC1.7 kinases are required for polar pollen tube growth. **A** Schematic representation of the proteins encoded by the Arabidopsis *PDK1* and *PDK2* genes and by the indicated splice variant *PDK1S0* or the mutant alleles *pdk1-b, pdk1-13, pdk1-14, pdk2-1* and *pdk2-4*. The protein serine/threonine kinase domain is depicted in orange and the PH domain in blue. **B** Quantification of the pollen tube phenotypes (normal: white bar; twisted: gray bar; swollen: black bar) observed in wild-type Arabidopsis (Col-0) or in the *pdk1-14 pdk2-1 p35S:YFP-PDK1, pdk1-b pdk2-1, pdk1-14 pdk2-1 pPDK1:YFP-PDK1, pdk1-14 pdk2-1 pPDK1:YFP-PDK1S0*, or *agc1*.*5 agc1*.*7* mutant lines. At least 300 pollen tubes (300 to 1000) for each line were quantified. Data were analyzed using two-way ANOVA followed by Tukey’s test. Letters indicate different significance groups (p < 0.0001, Error bar: standard error of mean). **C-H** Differential interference contrast microscopy images showing the pollen tube phenotypes of wild-type Arabidopsis (Col-0) in the *pdk1-14 pdk2-1 p35S:YFP-PDK1, pdk1-14 pdk2-1 pPDK1:YFP-*PDK1, pdk1*-b pdk2-1, pdk1-14 pdk2-1 pPDK1:YFP-PDK1S0* or *agc1*.*5 agc1*.*7* mutants lines. Red arrow indicates a swollen *pdk1-b pdk2-1* pollen tube tip. **I-J**, Fluorescence microscopy images of the pollen tubes of *pdk1 pdk2 pPDK1:YFP-PDK1* (**I**),; and *pdk1 pdk2 pPDK1:YFP-PDK1S0* (**J**). Inset in **I** shows a 5 times zoomed in image of different pollen tube of the same line. Please note that YFP:PDK1S0 cannot be detected in the pollen tube tip (red arrow) in **J**. Scale bars indicate 50 µm.

As a first approach to study this, we compared the pollen tube phenotype of wild-type (Col-0), *pdk1-b pdk2-1, pdk1 pdk2 pPDK1:YFP-PDK1S0, pdk1 pdk2 p35S:YFP-PDK1* and *pdk1 pdk2 pPDK1:YFP-PDK1* (Fig. 1B-H). The *pdk1 pdk2* lines were not used, as these plants grow very slow. Instead *pdk1 pdk2 p35S:YFP-PDK1* was used, as the *p35S:YFP-PDK1* construct rescues the vegetative defects but is not expressed in pollen (Wilkinson et al., 1997; Xiao and Offringa, 2020). Both wild-type and *pdk1 pdk2 pPDK1:YFP-PDK1* showed normal pollen tube growth, whereas *pdk1 pdk2 p35S:YFP-PDK1* pollen showed either twisted or curly tubes (here both types are referred to as “twisted”), or tubes with a swollen tip (referred to here as “swollen”) at approximately a 1:1 ratio (Fig. 1B, D). Generally, the swollen tubes were shorter, indicating that their growth had ceased. Interestingly, *pdk1-b pdk2-1* and *pdk1 pdk2 pPDK1:YFP-PDK1S0* both showed about 90% twisted tubes and 10% swollen tubes (Fig. 1B, F, G). This suggests that PDK1 has a dual role in pollen tube growth. When the PDK1 kinase domain is defective, growth is lost, resulting in short tubes with swollen tips. When PDK1 lacks the PH domain but retains kinase activity, growth persists but its polarity is lost, causing tube twisting. These results indicate that PDK1 promotes growth via its kinase domain and controls polarity via its PH domain. Consistently, YFP–PDK1 accumulated at pollen tube tips (Fig. 1I), whereas YFP– PDK1S0 was undetectable (Fig. 1J), likely reflecting rapid turnover of the PH domain–lacking protein.(Bayascas *et al*., 2008; Xiao and Offringa, 2020).

### AGC1.5 and AGC1.7 act redundantly downstream of PDK1 in regulating pollen tube growth

The *agc1*.*5 agc1*.*7* double mutant also shows defects in pollen tube growth (Zhang *et al*., 2009), with a 1:1 ratio of twisted or swollen tubes, similar to *pdk1 pdk2 p35S:YFP-PDK1* pollen (Fig. 1B, D, H). These comparable defects, along with reports that PDK1 phosphorylates AGC1.5 and AGC1.7 (Li *et al*., 2018; Zegzouti *et al*., 2006) suggested that AGC1.5/1.7 act redundantly downstream of PDK1.

Expression of a constitutively active version of AGC1.5 (AGC1.5SD) N-terminally fused with YFP under the *PDK1* promoter did not rescue pollen tube growth in *pdk1 pdk2* or *pdk1-b pdk2-1* mutants (Fig. 2A, B). As no fluorescent signal could be detected, we expressed the fusion protein under the pollen-specific *LAT52* promoter in the *pdk1-b pdk2-1*. We observed normal growth of the YFP positive pollen tubes in the T1 generation, whereas the YFP negative tubes retained mutant phenotypes (Fig. 2 C). Quantification showed that 95% of the YFP positive tubes showed normal growth, whereas all YFP-negative tubes showed defective growth (Fig. 2D). Interestingly, YFP-AGC1.5SD rescued polar growth without tip localization (Fig. 2C). Consistently, mCherry-AGC1.5 was localized throughout the cytosol in wild-type tubes, whereas YFP-PDK1 accumulated at the tip (Fig. 2E). These data indicate that polar PDK1 accumulation in wild-type pollen appears sufficient to locally activate AGC1.5/1.7 and maintain growth polarity. Pollen-specific expression of constitutively active AGC1.5SD driven by the *LAT52* promoter, but not the weaker *PDK1* promoter, is sufficient to activate tip-localized targets such as RopGEFs (Li *et al*., 2018), and support normal tube growth.

**Figure 2.**
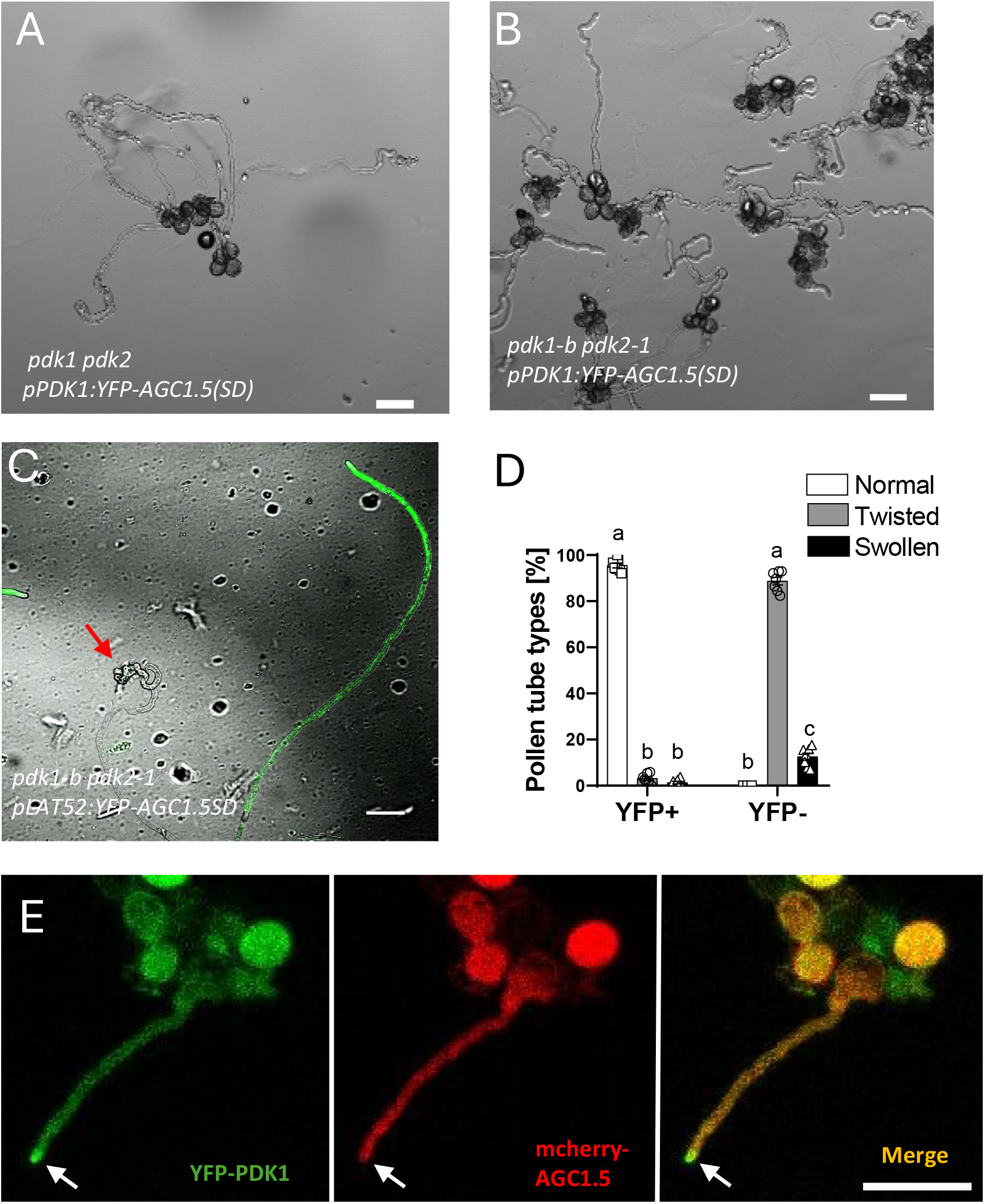
The aberrant pollen tube phenotype of *pdk1-b pdk2-1* are rescued by AGC1.5SD. **A-B** Differential interference contrast microscopy images showing the pollen tube phenotype of Arabidopsis lines *pdk1 pdk2 pPDK1:YFP-AGC1*.*5SD* (**A**) and *pdk1-b pdk2-1 pPDK1:YFP-AGC1*.*5SD* (**B**). **C** Fluorescent microscopy image showing the phenotypes of T1 *pdk1-b pdk2 pLAT52:YFP-AGC1*.*5SD* pollen tubes. The YFP-AGC1.5SD expressing pollen tube shows normal growth, whereas the red arrow indicates the twisted growth of a non-expressing pollen tube. **D** Quantification of the pollen tube phenotypes of the YFP-AGC1.5SD expressing (YFP+) and non-expressing (YFP-) pollen tubes in **C**. 418 pollen tubes from three different lines were quantified. Data were analyzed using two-way ANOVA followed by Tukey’s test. Letters indicate different significance groups (p < 0.0001, Error bar: standard error of mean). **E** Confocal microscopy images of germinating pollen of a *pdk1 pdk2 pPDK1:YFP-PDK1 pLAT52:mCherry-AGC1*.*5* T2 plant. White arrow indicates the pollen tube tip. Scale bars indicate 50 µm.

### PDK1 and PI(4,5)P_2_ maintain co-localization at the pollen tube tip through feedback regulation

The role of the PDK1 PH domain in pollen tube polarity prompted us to examine upstream phospholipids regulating PDK1. In animal cells, PI(3,4,5)P_3_ is the primary activator of PDK1 (Shaw and Burke, 2025), but this lipid is absent in plants (van Leeuwen et al., 2004), and the Arabidopsis PDK1 PH domain binds phospholipids promiscuously (Deak et al., 1999). We therefore focused on PI(4,5)P_2_, as it activates PDK1 *in vitro* (Anthony et al., 2004), localizes to the pollen tube tip (Potocký et al., 2014), and its biosynthesis enzymes (PIP5Ks) are required for pollen germination and tube growth (Ischebeck *et al*., 2008; Ischebeck *et al*., 2011; Kato et al., 2024; Sousa *et al*., 2008; Zhao *et al*., 2025).

Introducing the PI(4,5)P_2_ reporter *P24R* into the *pdk1 pdk2 pPDK1:YFP-PDK1* background revealed that PI(4,5)P_2_ accumulates at the pollen germination site before PDK1 (Supplementary Fig. 1A, 4 to 7 min), which appears about 2 min later at the center of the PI(4,5)P_2_ signal, strengthens within 3 minutes (Supplementary Fig.1A, 10 min), and localizes to the tip upon tube emergence (Supplementary Fig.1A, 12 min). Kymograph analysis and fluorescence quantification at the germination point (Supplementary Fig. 1A, 9 min, red arrow) further supported this. The PDK1 signal appeared at 9 min, whereas PI(4,5)P_2_ was already strongly enriched at the germination site at 4 min (Supplementary Fig. 1B), suggesting that the PI(4,5)P_2_ acts upstream to recruit and activate PDK1 at the germination site. During tube growth, PDK1 and PI(4,5)P_2_ co-localized at the tip, with PI(4,5)P_2_ forming a dome at the plasma membrane with PDK1 mainly cytosolic beneath it (Fig. 3A).

**Figure 3.**
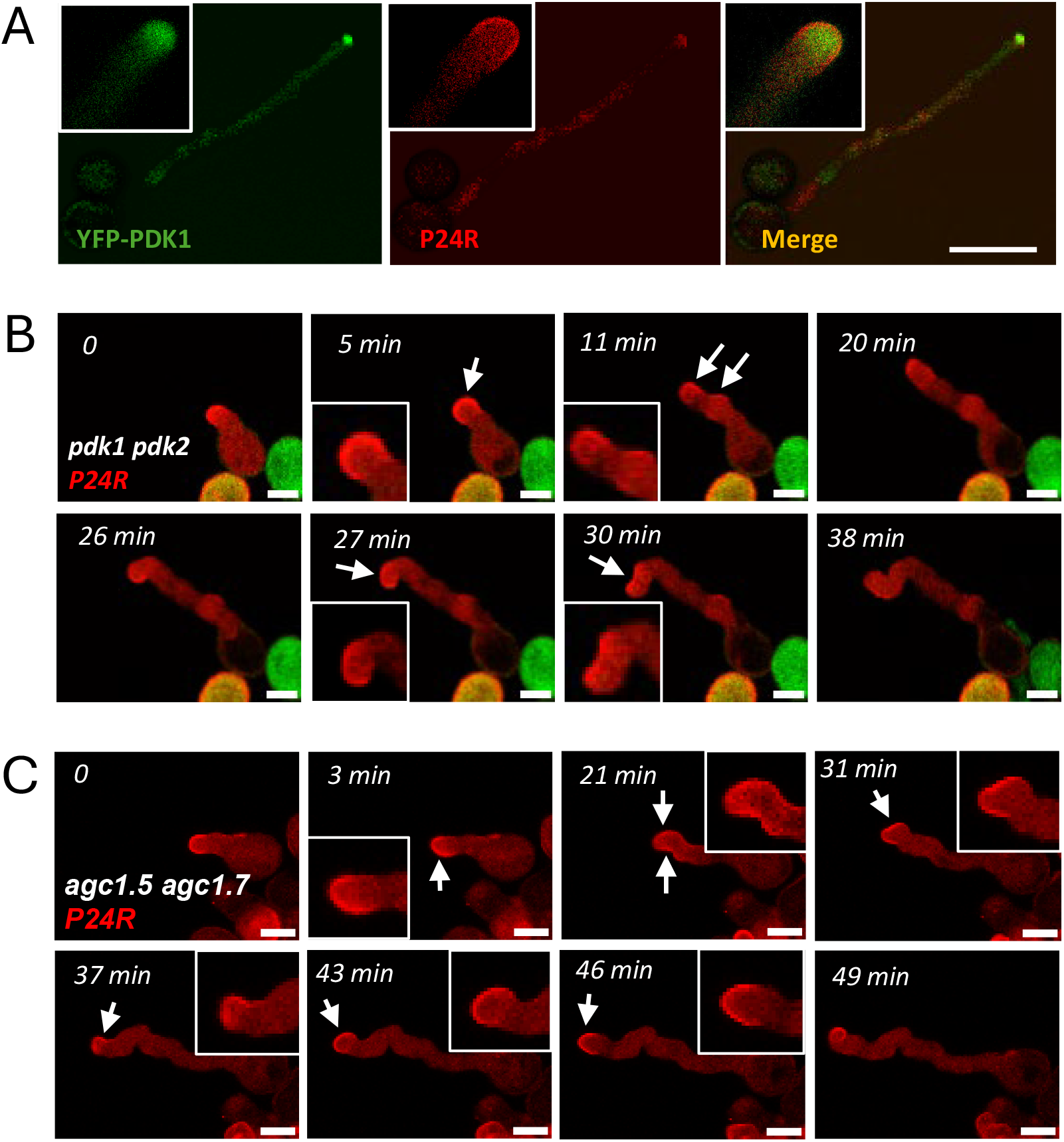
PI(4,5)P_2_ localization in wild type, *pdk1 pdk2* or *agc1*.*5 agc1*.*7* pollen tubes. **A** Confocal microscopy images showing the localization of YFP-PDK1 (left) and PI(4,5)P_2_ (P24R reporter, middle) in *pdk1 pdk2 pPDK1:YFP-PDK1 P24R* pollen tubes, scale bar indicates 50 µm. Images in upper-left corner show a zoomed in view of the pollen tube tip. **B, C** Confocal microscopy time-lapse images recording the pollen tube growth of *pdk1 pdk2 P24R* (**B**) or *agc1*.*5 agc1*.*7 P24R* (**C**) pollen. Pollen in (**B**) show segregation of the fluorescent marker as these are derived from a *pdk1 pdk2 pPDK1:YFP-PDK1* +/- *P24R* +/- plant. White arrows point at the wider more diffuse PI(4,5)P_2_ distribution. Scale bars indicate 10 µm. Images in the corner show a zoomed in view of the pollen tube tip.

Time-lapse imaging of *P24R* in *pdk1 pdk2* and *agc1*.*5 agc1*.*7* mutants showed that the symmetric PI(4,5)P_2_ dome at the tip is lost, often resulting in a wider or asymmetric distribution compared to wild type (Fig. 3B,C), which led to stalled growth and tip swelling (Fig. 3B 5-11min), or changes in growth direction (Fig. 3B 27-38 min, Fig. 3C 31-37 min). These data indicate that PI(4,5)P_2_ directs AGC kinase activity, which in turn feeds back to maintain the PI(4,5)P_2_ dome at the tube tip, ensuring polar pollen tube growth.

### PH domain restricts PDK1 and PI(4,5)P_2_ localization at the pollen tube tip

Five *PIP5K* genes function redundantly in Arabidopsis during pollen tube growth (Heilmann and Ischebeck, 2016; Ischebeck *et al*., 2008; Ischebeck *et al*., 2011; Kato *et al*., 2024; Sousa *et al*., 2008) and *pip5k4 pip5k5 pip5k6* triple mutant pollen do not germinate (Kato *et al*., 2024) prohibiting a mutant approach. Treatment with 0.5 or 1 μM UNC3230, a known PIP5K inhibitor in animal cells (Wright et al., 2014), strongly inhibited pollen germination and tube growth (Supplementary Fig. 2A,B), similar to what was observed for *pip5k* mutants. However, growth polarity and the localization of PDK1 or PI(4,5)P_2_ were not affected (Supplementary Fig. 2B-D), suggesting that reducing PI(4,5)P_2_ levels is not an appropriate way to study its function in polar tube growth.

Interestingly, treating with 10 or 50 μM PTH427, a compound competitively binding the PH domain of PDK1 or PKB in mouse cancer cells (Meuillet et al., 2010), resulted in short pollen tubes with a swollen tip, similar to *pdk1 pdk2* and *agc1*.*5 agc1*.*7* mutant tubes (Fig. 4A). In treated tubes, both YFP-PDK1 and P24R signals increased, with YFP-PDK1 signal no longer confined to the tip and an expanded P24R dome (Fig. 4B, C). The stronger yet dispersed YFP-PDK1 signal suggests that the inhibitor stabilizes the kinase while impairing its phospholipid-dependent tip recruitment. The broadened PI(4,5)P_2_ signal and swollen tips resemble those in *pdk1 pdk2* and *agc1*.*5 agc1*.*7* pollen tubes (Fig. 3B,C), supporting feedback between kinase activity and PI(4,5)P_2_ localization, focusing PDK1 at the tip to maintain polar growth.

**Figure 4.**
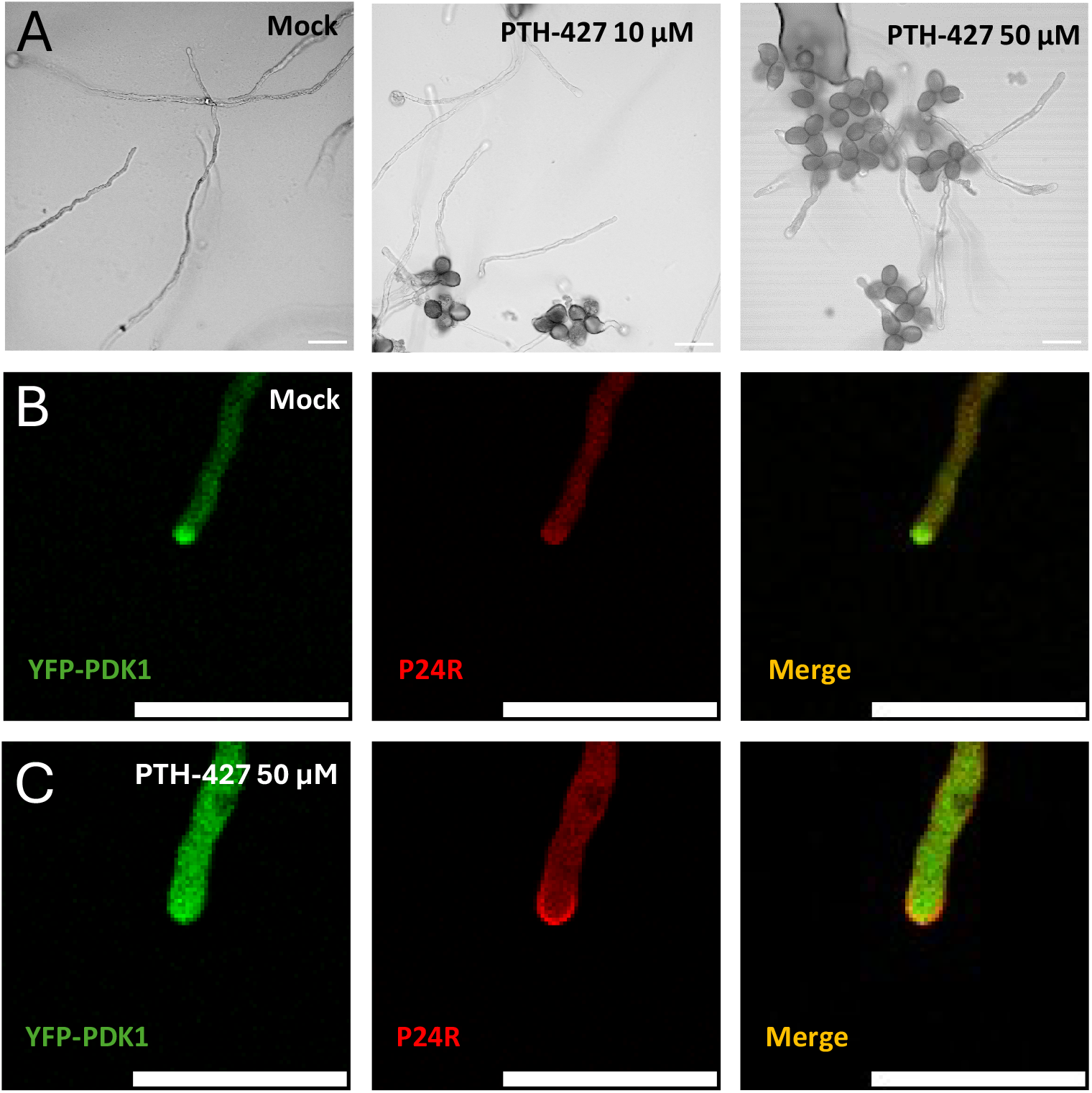
A PH domain inhibitor enhances PDK1 and PI(4,5)P_2_ abundance in pollen tube tips. **A** Brightfield microscopy images showing the pollen tube morphology of mock or PTH427 (10 and 50 µM) treated germinating *pdk1 pdk2 pPDK1:YFP-PDK1 P24R* pollen. **B, C** Confocal microscopy images showing the localization of YFP-PDK1 (left) and PI(4,5)P_2_ (P24R reporter, middle) in the pollen tubes of germinating *pdk1 pdk2 pPDK1:YFP-PDK1 P24R* pollen following mock (**B**) or 50 µM PTH427 (**C**) treatment. Scale bars indicate 50 µm.

### The PDK1-AGC1.5/1.7 kinase cascade is required for proper endocytosis and actin distribution

Endocytosis and actin organization are essential for pollen tube growth (Cheung *et al*., 2010; Guo and Yang, 2020; Vidali et al., 2001; Ye *et al*., 2009). Given that AGC1.5/1.7 act downstream of PDK1 and that pdk1 pdk2 and agc1.5 agc1.7 mutants show similar tube phenotypes, we examined endocytosis and actin organization in these backgrounds.

In wild-type pollen tubes stained with FM5-95, endocytosis was centered at the tube tip (Supplementary Fig. 3A). In twisted *pdk1 pdk2 pPDK1:YFP-PDK1S0* pollen tubes, endocytosis initially localized at the tip center but shifted to the side toward which the tube would curl before bending (Supplementary Fig. 3B, 0-2.5min). In short swollen *pdk1 pdk2 p35S:YFP-PDK1* tubes, endocytosis was initially centered but disappeared upon growth arrest (Supplementary Fig. 3C). This is consistent with our hypothesis that the PDK1 kinase domain is responsible for tube growth. Similar endocytic defects were observed in twisted or swollen *agc1*.*5 agc1*.*7* pollen tubes (Supplementary Fig. 3D, E).

Actin distribution in pollen tubes was visualized using phalloidin staining. In twisted mutant tubes, actin distribution resembled that of wild-type tubes as long as tubes grew straight, showing a normal collar and tip clear zone (Supplementary Fig. 4A). Upon twisting, actin cables followed the tube shape extending into bulges or branches (Supplementary Fig. 4B). In the swollen tubes, the cable actin protruded to and accumulated in the tip (Supplementary Fig. 4C). Similar actin defects in *pdk1 pdk2* and *agc1*.*5 agc1*.*7* pollen tubes further support that AGC1.5/1.7 act downstream of PDK1 and that this pathway regulates pollen tube growth via endocytosis and actin organization.

## DISCUSSION

Decades of research have identified multiple signaling pathways and cellular processes regulating polarized pollen tube growth, including endocytosis, actin organization and phosphoinositide-, AGC kinase- and ROP-mediated signaling (Fu, 2015; Guo and Yang, 2020; Heilmann and Ischebeck, 2016; Ou and Yi, 2022). However, how these pathways and cellular processes are integrated has remained unclear.

Here we show that the AGC kinases AGC1.5/1.7 act downstream of PDK1 to control pollen tube polarity and that PDK1, through its PH domain, is likely recruited by PI(4,5)P_2_ to the germination site. After germination, PDK1 and PI(4,5)P_2_ remain colocalized at the pollen tube tip through reciprocal feedback, with PI(4,5)P_2_ forming a dome at the tip membrane and PDK1 accumulating in the underlying cytosol. This PI(4,5)P_2_-PDK1-AGC1.5/1.7 cascade ensures polar growth by positioning endocytosis at the tip center, likely through modulating actin organization.

In animals, PH domain–mediated phospholipid binding is crucial for PDK1 function, and PH domain mutations severely impair multiple cellular processes (Bayascas *et al*., 2008). By contrast, in plants the PH domain appears largely dispensable for development, as the PH domain-lacking splice variant PDK1S0 rescues the growth and most developmental defects of *pdk1 pdk2* mutants (Xiao and Offringa, 2020). Whether the PH domain becomes important under abiotic or biotic stress remains unclear. For now, only polar pollen tube growth seems to require full length PDK1.

In animal cells, PI(3,4,5)P_3_ is the main target for the PDK1 PH domain causing its plasma membrane recruitment and activation (Shaw and Burke, 2025). Although the PDK1 PH domain can bind PI(3,4,5)P_3_ *in vitro* (Deak *et al*., 1999), PI(3,4,5)P_3_ is not found in plants (van Leeuwen *et al*., 2004). Instead, this PH domain binds several other phospholipids (Deak *et al*., 1999), among which phosphatidic acid (PA) and PI(4,5)P_2_ - but not PI(3)P or PI(4)P - enhance its kinase activity (Anthony *et al*., 2004). Of these, only PI(4,5)P_2_ localizes at the pollen tube tip (Potocký *et al*., 2014; Zhao *et al*., 2025). Moreover, the five pollen-expressed PIP5Ks are also tip-localized and essential for pollen germination and tube growth (Ischebeck *et al*., 2008; Ischebeck *et al*., 2011; Sousa *et al*., 2008; Zhao *et al*., 2025). Together with our observations on PDK1 recruitment to the germination initiation site and the feedback regulation between PI(4,5)P_2_ and the PDK1-AGC1.5/1.7 cascade at the tube tip, these data strongly suggest that PI(4,5)P_2_ is the key upstream regulator of PDK1 during pollen tube growth.

The AGC1.5 kinase and its constitutively active form AGC1.5SD are detected throughout the cytosol of the pollen tube, while PDK1 accumulates at the tip (Fig. 2E). In wild-type plants, tip-localized PDK1 locally activates the cytosolic AGC1.5/1.7 to phosphorylate downstream targets, such as RopGEFs (Li *et al*., 2018). As these RopGEFs are tip localized as well (Li *et al*., 2018), this may explain why cytosolic AGC1.5SD can still rescue the *pdk1 pdk2* phenotype, and why this requires its expression from the stronger pollen-specific *LAT52* promoter instead of the *PDK1* promoter.

Our results also indicate that the twisted pollen tube phenotype of *pdk1 pdk2* is due to an unstable endocytosis polarity, with the varying position of the endocytosis maximum directing twisted pollen tube growth, and with loss of endocytosis leading to swollen tube tips. This supports our current model that PDK1 through its PH domain maintains endocytic polarity via reciprocal feedback with PI(4,5)P_2_, whereas the kinase domain promotes growth. Recent work showed that oscillatory PI(4,5)P_2_ at the germination site is required for polarity establishment and pollen germination (Zhao *et al*., 2025), which is consistent with our observations on pre-germination PI(4,5)P_2_ oscillation (Supplementary Fig. 1B) and the strongly reduced germination upon PIP5K inhibitor treatment (Supplementary Fig. 2A). The PI(4,5)P_2_ oscillation is regulated by actin nucleation factor AtFH5-mediated exocytosis (Zhao *et al*., 2025), and seems required for assemble the polarity machinery before recruiting PDK1. Polar pollen tube growth requires a balanced exo- and endocytosis where PI(4,5)P_2_ contributes to endocytosis, as *PIP5K6* downregulation reduces endocytic activity (Zhao *et al*., 2010). Together, these observations highlight a coordinated role for PI(4,5)P_2_ and PDK1 in coupling membrane trafficking to polarity maintenance.

In germinating *pdk1 pdk2* or *acg1*.*5 agc1*.*7* pollen, twisted tubes with functional endocytosis show a relatively normal actin organization, whereas swollen tubes lacking endocytosis exhibit abnormal actin protrusions in the tip, suggesting that the PDK1-AGC1.5/1.7 signaling regulates endocytosis by modulating actin. Interestingly, the actin phenotype in twisted *pdk1 pdk2* tubes resembles that induced by *PIP5K4* or *PIP5K5* overexpression (Ischebeck *et al*., 2008), supporting a reciprocal regulatory loop between PDK1 and PI(4,5)P_2_ in maintaining actin organization and stable polar endocytosis.

ROP signaling controls actin-dependent endo- and exocytosis (Lee et al., 2008) and is activated by RopGEFs following phosphorylation by AGC1.5 (Li *et al*., 2018). Recent work showed that AGC1.7 phosphorylates ADF7 to regulate actin organization in pollen tubes (Li et al., 2024). Together with our findings, this leads to a model where pollen tube polarity is established by reciprocal feedback between PDK1 and PI(4,5)P_2_, the activated PDK1 phosphorylates and activates AGC1.5/1.7, these AGC kinases in turn phosphorylate ROPGEFs, and ROPGEFs subsequently regulate the ROP-dependent actin organization and endo/exocytosis. An additional branch in this model is that AGC1.5/1.7 directly regulate actin organization through ADF7.

## METHODS

### Plant lines, transformation procedures and growth conditions

*Arabidopsis thaliana* ecotype Columbia 0 (Col-0) was used as wild-type control. The mutant lines *pdk1-13 pdk2-4 pPDK1:YFP-PDK1, pdk1-13 pdk2-4 pPDK1:YFP-P1S0, pdk1-13 pdk2-4 p35S:YFP-PDK1* and *pdk1-14 pdk2-4* (Xiao and Offringa, 2020), *pdk1-b pdk2-1* (Camehl *et al*., 2011), and PI(4,5)P_2_ reporter line *P24R* (two rat PLCδ1 PH domains fused to Cherry and expressed under the *UBIQUITIN10* promoter (Simon et al., 2014)) are all in Col-0 background and have been described previously.

For transgenic lines created in this study, the T-DNA construct *pPDK1:YFP-AGC1*.*5SD* was transformed into *pdk1-14(-/-) pdk2-4(+/-)* and *pdk1-b pdk2-1*, and T3 homozygotes were used for analysis. The T-DNA constructs *pPDK1:YFP-AGC1*.*5SD-SPH, pPDK1:YFP-AGC1*.*5SD-LPH* and *pLAT52: YFP-AGC1*.*5SD* were transformed into *pdk1-b pdk2-1*, and T3 homozygotes were used for analysis. The T-DNA construct *pLAT52:mCherry-AGC1*.*5* was transformed into *pdk1-13 pdk2-4 pPDK1:YFP-PDK1*, and T1 generation plants were used for analysis. Binary vectors carrying these T-DNAs were introduced into *Agrobacterium tumefaciens AGL1* by electroporation (Den Dulk-Ras and Hooykaas, 1995), correct strains were selected on Luria-Bertani medium plates containing 75 ng/L carbenicillin (for agrobacteria) and 100 ng/L kanamycin (for the transformed construct), and plant transformation was performed using the floral dip method (Clough and Bent, 1998). Transgenic plants were selected using 50 mg/L kanamycin. In case of transformations in the *pdk1-14(-/-) pdk2-4(+/-)* background, the T1 seedlings were genotypes for homozygosity of the *pdk2-4* allele. The *pdk1 pdk2 pPDK1:YFP-PDK1* line was crossed with the PI(4,5)P_2_ reporter line *P24R*, and pollen of T2 generation plants were used for the analysis. The *agc1*.*5 agc1*.*7* double mutant was obtained by crossing the *agc1*.*5* (*SALK_073610C*) and *agc1*.*7-1* (*SALK_140378C*) alleles. Dreamtaq DNA polymerase (ThermoFisher, EP0705, Waltham, USA) was used for genotyping using primers listed in Table S1.

For seedling growth, seeds were surface-sterilized by 1 minute in 70% ethanol, 10 minutes in 1% chlorine followed by five washes with sterile water. Sterilized seeds were kept in the dark at 4°C for 2 days for vernalization and germinated on vertical plates with 0.5× Murashige and Skoog (1/2 MS) medium (Duchefa, M0222, Haarlem, the Netherlands) containing 0.05% MES, 0.8% agar (Daishin, D1004, Haarlem, the Netherlands) and 1% sucrose at 22 °C and 16 hours photoperiod. Plants were grown on soil at 21°C, 16 hours photoperiod, and 70% relative humidity.

### Cloning procedures

To obtain *pLAT52: YFP-AGC1*.*5SD, AGC1*.5 was PCR amplified from Col-0 cDNA using primers listed in Table S1 and cloned into *pDONR207* by a Gateway BP reaction (Invitrogen, Gateway BP Enzyme Mix #11789020, Waltham, USA). The resulting plasmid *pDONR207-AGC1*.5 was used for QuickChange II XL Site-Directed Mutagenesis (Agilent, #200521, Santa Clara, the United States) with the primers listed in Table S1, to substitute serine (S) 408 in AGC1.5 for aspartic acid (D). The resulting *AGC1*.*5SD* coding region was then introduced into *pLAT52::YFP-gateway* by LR reaction (Invitrogen, Gateway LR Clonase II Enzyme Mix #12538120, Waltham, USA). *pLAT52::YFP-gateway* was generated by replacing the promoter of *pPDK1::YFP-gateway* (Xiao and Offringa, 2020). For *pPDK1:YFP-AGC1*.*5SD*, the *AGC1*.*5SD* coding region was introduced by LR reaction from *pDONR207-AGC1*.*5SD* into *pPDK1:YFP-gateway* (Xiao and Offringa, 2020). For *pLAT52:mCherry-gateway*, the segment of *Kpn*I-*mcherry*-*Xho*I was PCR amplified from *pART7-Gateway-mcherry* (Gleave, 1992) excised with *Kpn*I and *Xho*I, and subsequently ligated into *pLAT52:YFP-gateway* digested with *Kpn*I and *Xho*I, replacing the *YFP* coding region. The *pLAT52:mCherry-AGC1*.*5* was obtained by LR reaction with *pDONR207-AGC1*.*5*. Phusion™ High-Fidelity DNA Polymerase (ThermoFisher, F530S, Waltham, USA) was used for all PCR reactions. All new clones were verified by sequencing.

### Pollen culturing, pharmacology and microscopy

*In vitro* pollen culture was performed as described previously (Boavida and McCormick, 2007). In short, warm liquid pollen germination medium (PGM) containing 10 % Sucrose (w/v), 0.01 % Boric Acid (w/v), 5 mM KCl, 5 mM CaCl_2_, 1 mM MgSO_4_ and 1.5 % Agarose (w/v), pH 7.5 (with 100 mM KOH) was pipetted onto a microscopy slide in thin pads of approximately 1cm^2^. Arabidopsis pollen were placed on the solidified PGM pads and for pollen germination the slides were incubated at 28°C for 2.5h placed in a 20 cm-diameter petri dish with a moist tissue at the bottom and the lid closed. Following incubation the medium pads were covered with cover glass and pollen tube phenotypes were examined using a ZEISS Imager.M2 microscope (Jena, Germany)equipped with a ZEISS Axiocam 712 color camera. For time-lapse imaging, after applying pollen on the medium pads, a cover slip was put on top and the space between the cover slip and the pad was sealed with paraffine oil to protect from drying. Time-lapse images were taken with a Nikon AX confocal microscope (Tokyo, Japan) equipped with YFP and RFP laser, NSPARC detector and an incubator space with the temperature set at 28°C. Different areas on the PGM pad were selected using the microscope software and time lapse was performed with 1 min time interval.

For PTH-427 and UNC3230 treatment, 50 µM PTH-427 (MedChemExpress, HY-12063, New Jersey, USA) or 0,1/0,5/1 µM UNC3230 (MedChemExpress, HY-110150, New Jersey, USA) was directly added to the liquid PGM at the indicated concentration, and pads with pollen added were incubated 2h before imaging.

Acting was stained with Alexa Fluor 488 phalloidin (ThermoFisher, A12379, Waltham, USA) as described (Qu et al., 2020). For endocytosis staining, 5 µM FM5-95 (ThermoFisher, T23360, Waltham, USA) was added to the PGM. The endocytosis process was recorded by time-lapse imaging using Nikon AX confocal microscope (Tokyo, Japan) as described above.

## AUTHOR CONTRIBUTIONS

RO conceived the project and provided additional funding, TZ conducted the experiments, TZ and RO designed the experiments and analyzed the data, TZ wrote the original draft which was reviewed and edited by RO.

## ACKNOWLEDGMENTS

We thank Gerda Lamers and Joost Willemse for support with microscopy and Altay Temel for help with plant growth. We thank Prof. Yvon Jaillais for sharing the *P24R* line.

## DECLARATION OF INTERESTS

The authors declare no competing interests.

**Table S1.**
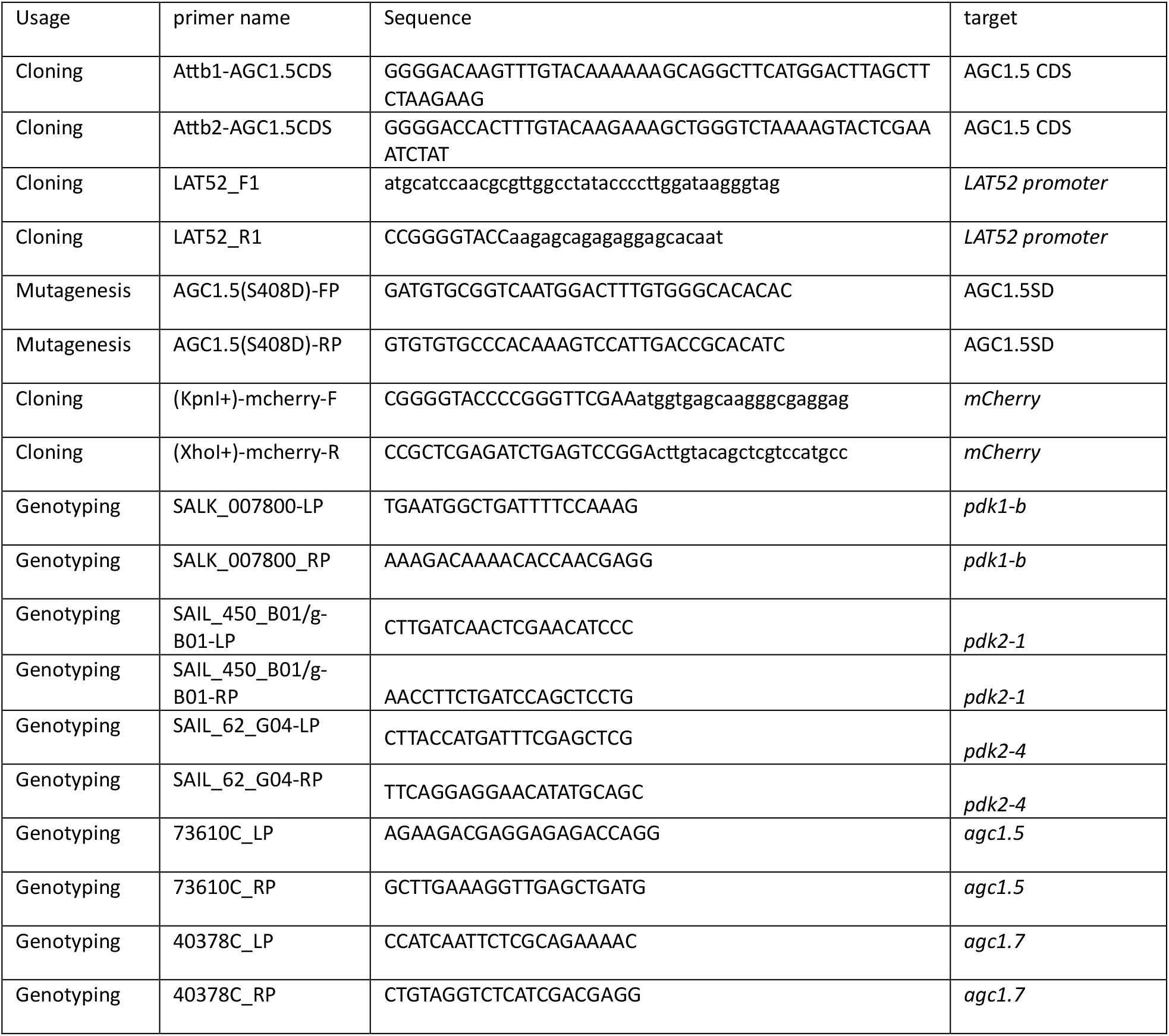
Primers used in this study.

## Figure legends

**Supplementary Figure 1.**
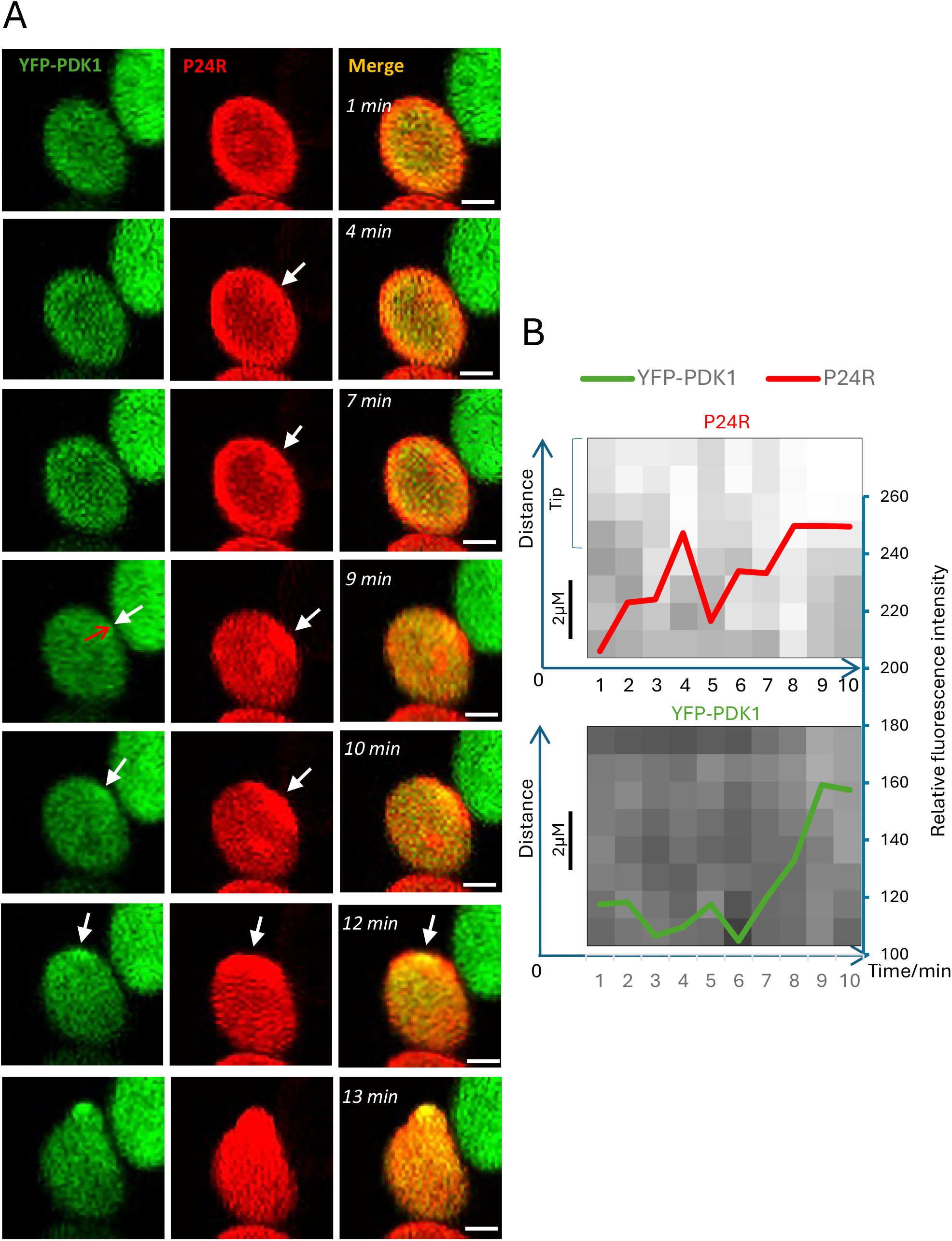
PI(4,5)P_2_ accumulates at the pollen germination site before PDK1. **A** Confocal microscopy time-lapse imaging over a 13-minute timeframe showing YFP-PDK1 (left) and PI(4,5)P_2_ (P24R reporter, middle) localization during the germination initiation of a *pdk1 pdk2 pPDK1:YFP-PDK1 P24R* pollen. The white arrow indicates the position of the germination initiation site. The red arrow at 9 min indicates the line used for kymograph analysis. Scale bars indicate 10 µm. **B** Kymograph analysis of the YFP-PDK1 and P24R signal from minute 1 to minute 10 along the red arrow shown in the YFP-PDK1 9 min image in **A**,. Each square represents one pixel, the shading inversely reflects the fluorescence intensity. The line graph depicts the mean relative fluorescence intensity of 4 pixels at the tip of the arrow.

**Supplementary Figure 2.**
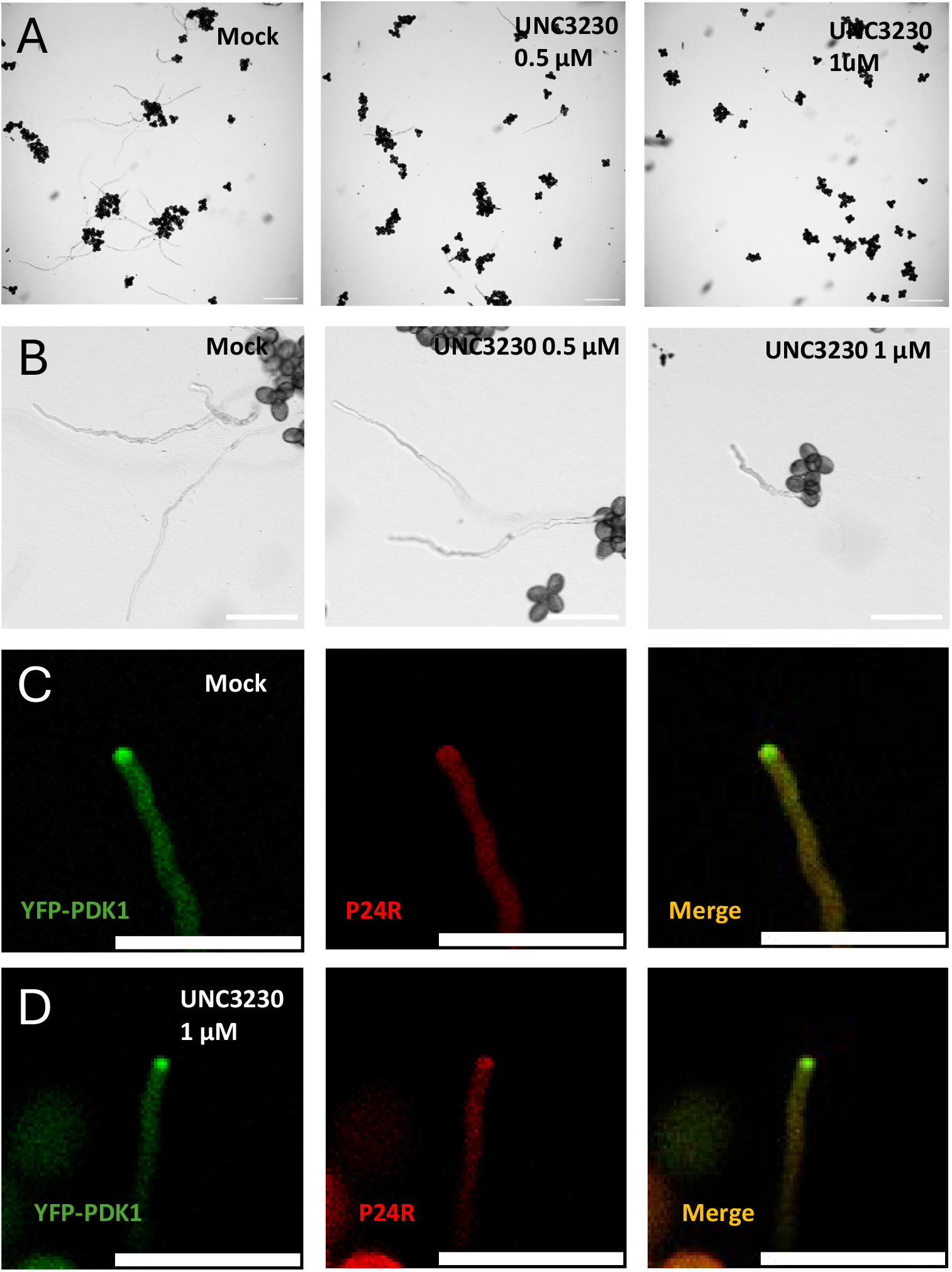
The PIP5K inhibitor UNC3230 suppresses pollen germination. **A, B** Brightfield microscopy images showing the germination rate (**A**) and pollen tube morphology (**B**) of mock or UNC3230 (0.5 µM and 1 µM) treated *pdk1 pdk2 pPDK1:YFP-PDK1 P24R* pollen. Scale bars indicate 200 µm. **C, D** Confocal microscopy images showing the localization of YFP-PDK1 (left) or PI(4,5)P_2_ (P24R reporter, middle) in mock (**C**) or 1 µM UNC3230 (**D**) treated *pdk1 pdk2 pPDK1:YFP-PDK1 P24R* pollen tubes. Scale bars indicate 50 µm.

**Supplementary Figure 3.**
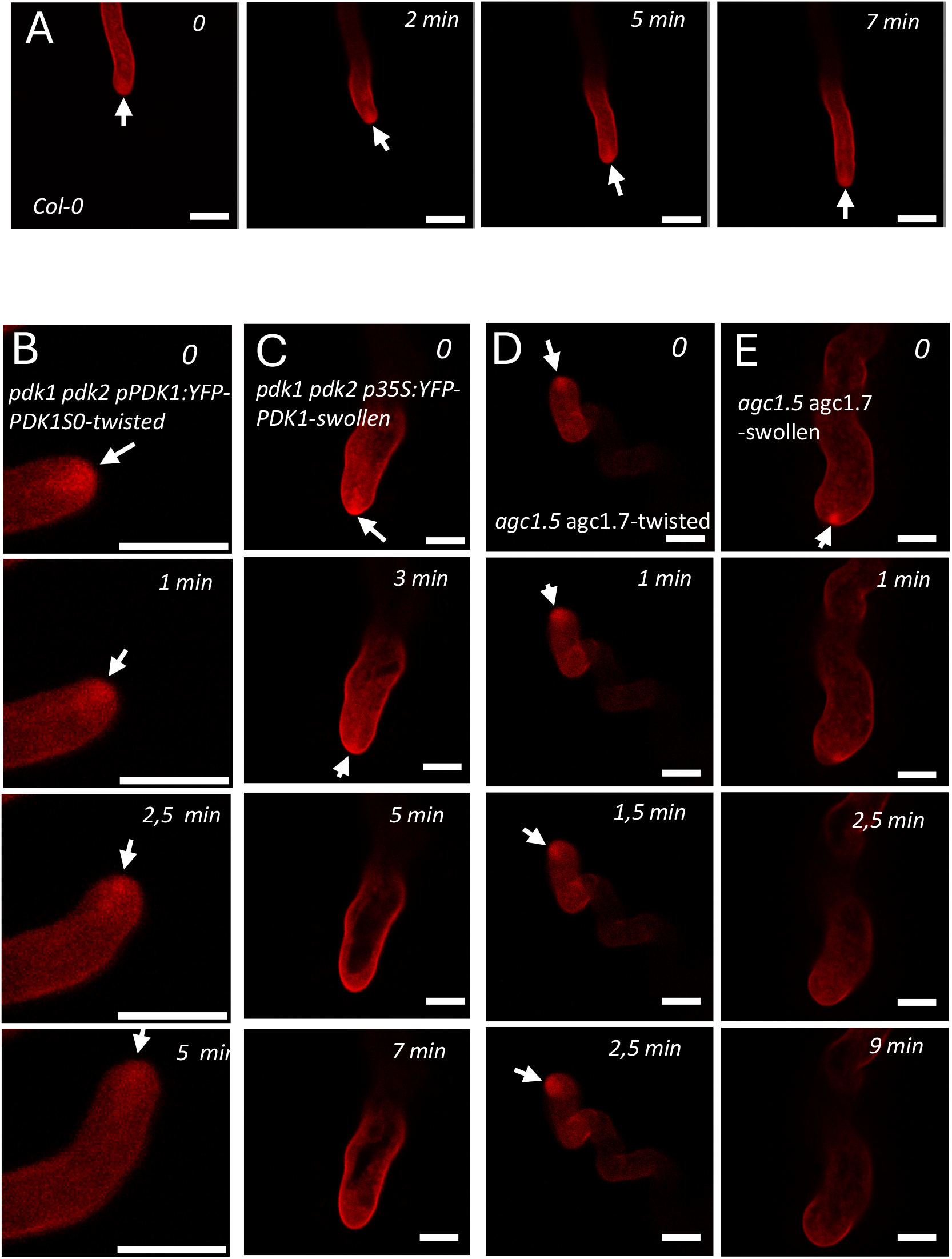
*pdk1 pdk2* and *agc1*.*5 agc1*.*7* pollen tubes show similar defects in endocytosis. **A-E** Confocal microscopy time-lapse images showing endocytosis in FM5-95 stained wild-type (Col-0) (**A**) twisting *pdk1 pdk2 pPDK1:YFP-PDK1S0* (**B**) swollen *pdk1 pdk2 pPDK1:YFP-PDK1S0* (**C**) twisting *agc1*.*5 agc1*.*7* (**D**) or swollen *agc1*.*5 agc1*.*7* (**E**) pollen tubes. Notes: Figure **B** shows zoomed-in images of a twisted pollen tube tip where the endocytosis is asymmetrically positioned to the side of pollen tube growth. Scale bar indicates 10 µm.

**Supplementary Figure 4.**
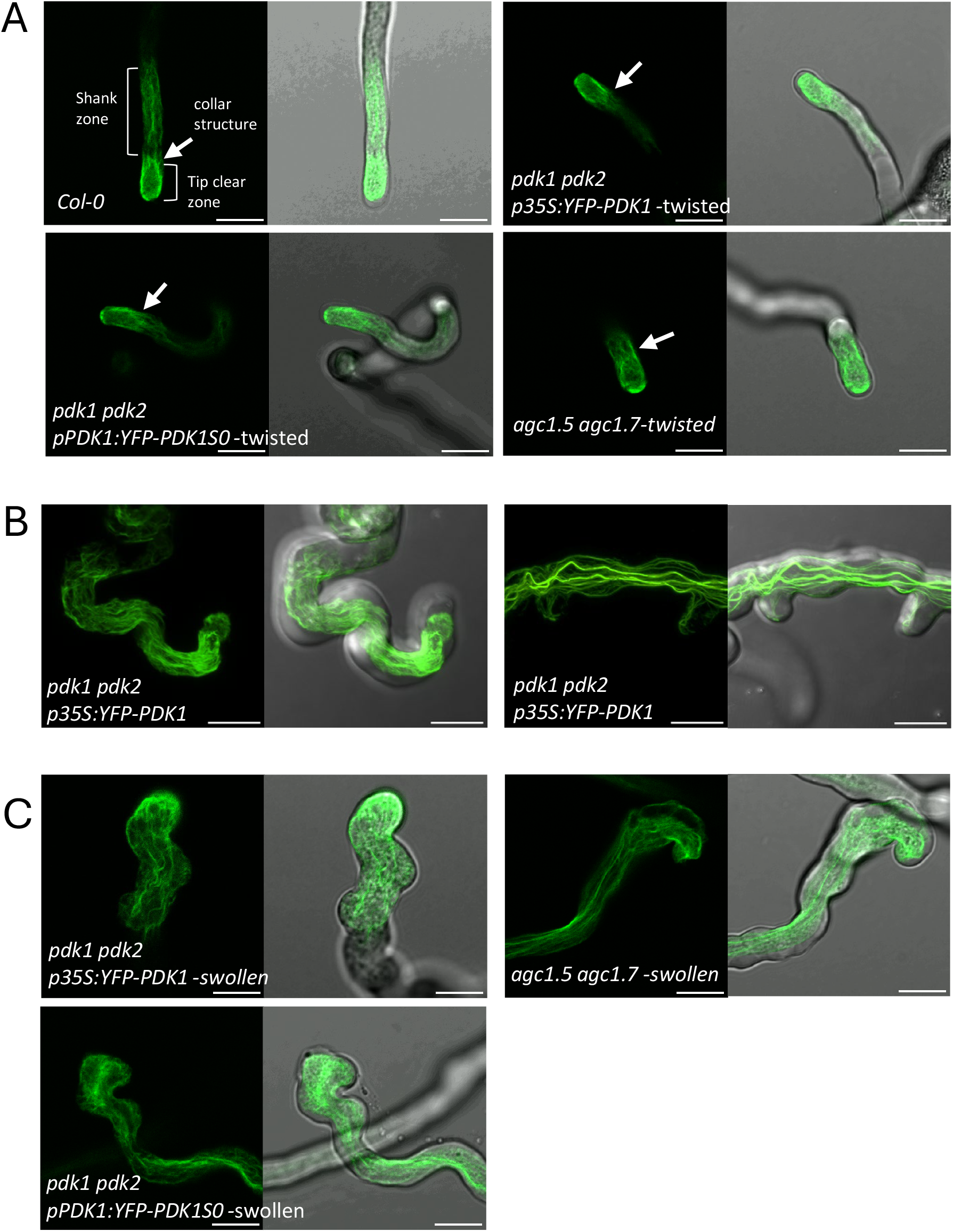
*pdk1 pdk2* and *agc1*.*5 agc1*.*7* pollen tubes show similar defects in actin distribution. **A-C** Confocal microscopy images of phalloidin 488**-**stained pollen tubes, showing the actin distribution in the tip part of wild-type (Col-0) or twisted *pdk1 pdk2 p35S:PDK1S0* or *agc1*.*5 agc1*.*7* pollen tubes (**A**), or in the twisted part of *pdk1 pdk2 p35S:PDK1S0* pollen tubes (**B**), or in the swollen tip of *pdk1 pdk2 p35S:PDK1S0* or *agc1*.*5 agc1*.*7* pollen tubes (**C**). Scale bar indicates 50 µm.

